# Evolution of sex determination and heterogamety changes in section *Otites* of the genus *Silene*

**DOI:** 10.1101/325068

**Authors:** Veronika Balounova,, Roman Gogela, Radim Cegan, Patrik Cangren, Jitka Zluvova, Jan Safar, Viera Kovacova, Roberta Bergero, Roman Hobza, Boris Vyskot, Bengt Oxelman, Deborah Charlesworth, Bohuslav Janousek

## Abstract

Switches in heterogamety occasionally occur both in animals and plants, although plant sex determination systems are mostly more recently evolved than those of animals, and have had less time for switches to occur. However, our previous research revealed a switch in heterogamety in section *Otites* of the plant genus *Silene*.

Here we analyse in detail the evolution of genetic sex determination in section *Otites*, which is estimated to have evolved about 0.55 MYA. Our study confirms female heterogamety in *S. otites* and newly reveals female heterogamety in *S. borysthenica*. Sequence analyses and genetic mapping show that the sex-linked regions of these two species are the same, but the region in *S. colpophylla*, a close relative with male heterogamety, is different. The sex chromosome pairs of *S. colpophylla* and *S. otites* each correspond to an autosome of the other species, and both differ from the XY pair in *S. latifolia*, in a different section of the genus. Our phylogenetic analysis suggests a possible change from female to male heterogamety within *Silene* section *Otites,* making these species suitable for detailed studies of the events involved.

## Introduction

Genetic sex determining systems in plants with separate sexes (dioecious plants) are thought mostly to have evolved via gynodioecy, involving a male sterility mutation in a hermaphroditic or monoecious ancestor as a first step^1^, or sometimes gradually from a monoecious ancestor, referred to as the paradioecy pathway^2^–^8^. The gynodioecy route is predicted to generate male heterogamety more often than female heterogamety, because male sterility mutations are mostly recessive, consistent with females being the homozygous sex^1^. It has therefore been suggested that species with female heterogamety evolved from ancestral systems with male heterogamety. However, evidence on the directions of changes in heterogamety (which are known in several taxa) is not yet sufficient to conclude that this is true. Moreover, some species with female heterogamety appear to have evolved without a previous male heterogametic system, for example in the genus *Fragari*a (*Rosaceae*). This well-studied genus includes gynodioecious and subdioecious species, and dioecious species. The subdioecious and dioecious species have female heterogamety, with dominant male sterility alleles in *F. virginiana* ^9,10^ and *F. chiloensis*^11^. In two other genera where multiple species have been studied, *Cotula*^3^ (*Asteraceae*), and *Pistacia*^12,13^ (*Anacardiaceae*), only female heterogamety has been found, again suggesting that there was no previous history of male heterogamety. In two further genera with female heterogamety, only single species have been tested (*Distichlis*^14^ in the *Poaceae*, and *Potentilla*^15^ in the *Rosaceae*).

Only two plant genera have so far been described as having both male and female heterogamety, *Populus*^16^, in the family *Salicaceae*, and *Silene*^17,18^, in the family *Caryophyllaceae*. In the *Salicaceae*, almost all species are dioecious, and the available data on fossils suggest that sex determination may be as old as 45 My^19,20^ so it is difficult to infer the ancestral system, as multiple changes could have occurred during this family’s long evolutionary history. It, therefore, remains unclear whether the different heterogametic states represent switches in heterogamety, and (if so) in which direction, or independent evolution of dioecy in different lineages^21^–^24^.

Dioecy in the *Caryophyllaceae* probably evolved much more recently than in *Populus*, probably within the past 10 million years^18,25^–^27^, making multiple changes in sex determination less likely than in the Salicaceae. In the genus *Silene*, at least three independent origins of genetic sex determination (dioecy and/or subdioecy) have nevertheless been reported ^18^, and a switch from XY to ZW, or vice versa, clearly occurred in section *Otites*^18^. In the present study, we identified sex-linked genes in *S. borysthenica* (section *Otites*, group *Cyri*), and *S. colpophylla* and *S. otites* (section *Otites*, group *Otites*), allowing us to test whether male heterogamety was ancestral in the section *Otites* and ancestral for all three group *Otites* species (the two species just mentioned, and *S. pseudotites*, see Fig. 1), and also whether the chromosomes carrying the sex-determining loci of different dioecious species evolved from the same autosome pair. The results allow us to exclude some of the theoretically possible models for evolutionary changes in heterogamety among these plants. We also discuss how female heterogamety might evolve de novo from an initially hermaphroditic species, via gynodioecy.

**Figure 1.**
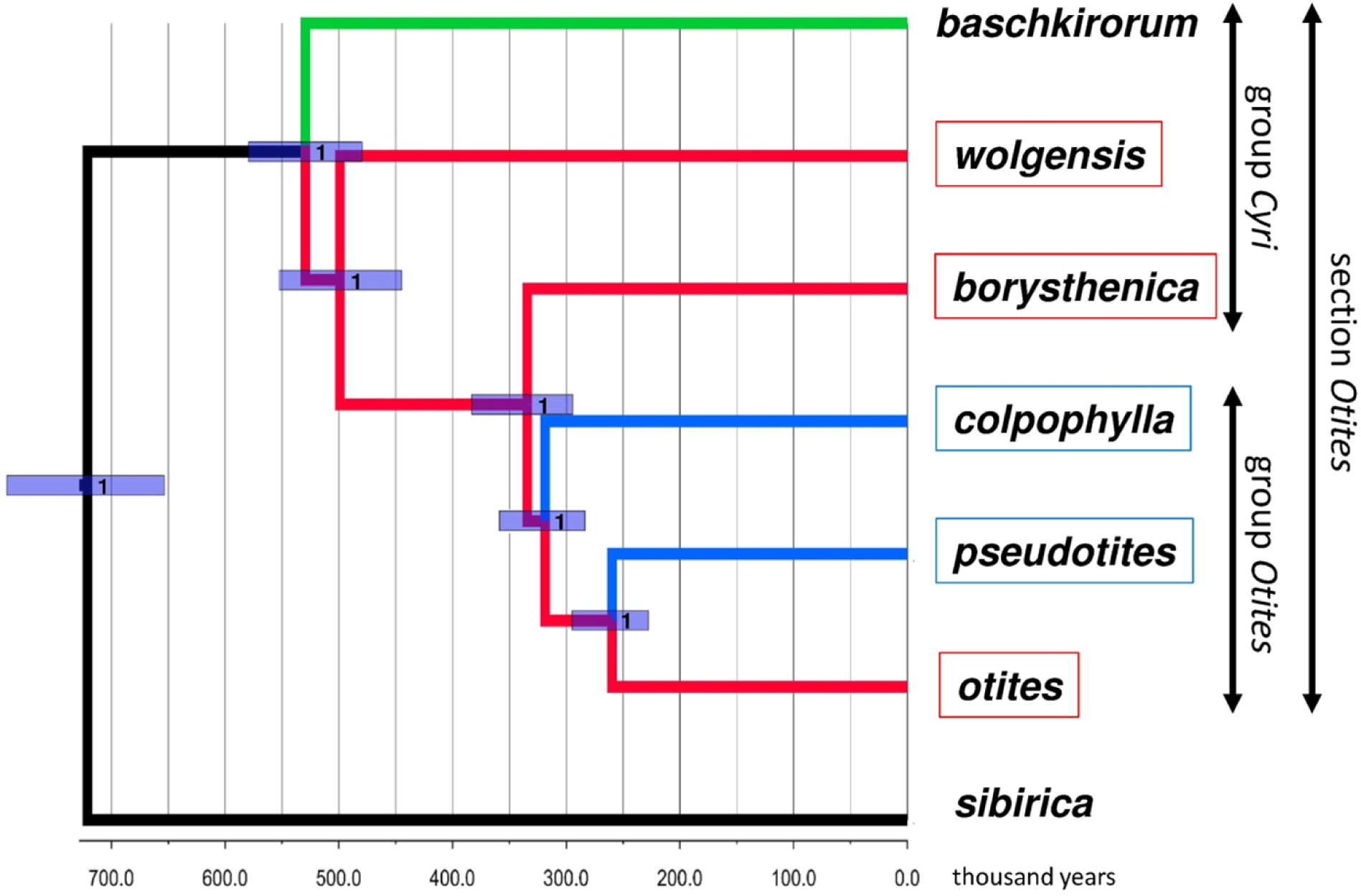
Results of StarBeast phylogenetic analysis of 662 single-copy orthogroup sequences. Species with female heterogamety are indicated by red boxes and male heterogamety by blue boxes. Female heterogamety in *Silene borysthenica*, and *S. otites*, and male heterogamety in *S. colpophylla*, were confirmed or inferred from the molecular data presented here, and the male heterogamety in *S. pseudotites* and female heterogamety in *S. wolgensis* from results of interspecific crosses^17,34^ (Newton, 1931 and Sansome, 1938). The values indicated at the nodes are posterior probabilities. Branch colours indicate the most probable scenario for the evolution of the sex determining system in section *Otites*. Branches and nodes where female heterogamety is inferred are coloured red, while the branches with male heterogamety are in blue. The branch to *S. baschkirorum* is coloured green as its heterogametic state is not known. *Silene sibirica* is not dioecious.

## Results

### Illumina RNAseq sequencing, 454 RNAseq sequencing and genetic analyses

The overall results of the different approaches employed in this study suggest that sex determining regions are found on three different chromosomes in the genus *Silene*: two different *S. latifolia* autosomes carry the *S. otites* and *S. borysthenica* (W) and *S. colpophylla* (Y) sex determining regions, and neither is homologous with the *S. latifolia* XY pair (Fig. 2A). We next describe the evidence that supports these conclusions, which are based on new, detailed studies of three *Otites* section species.

**Figure 2.**
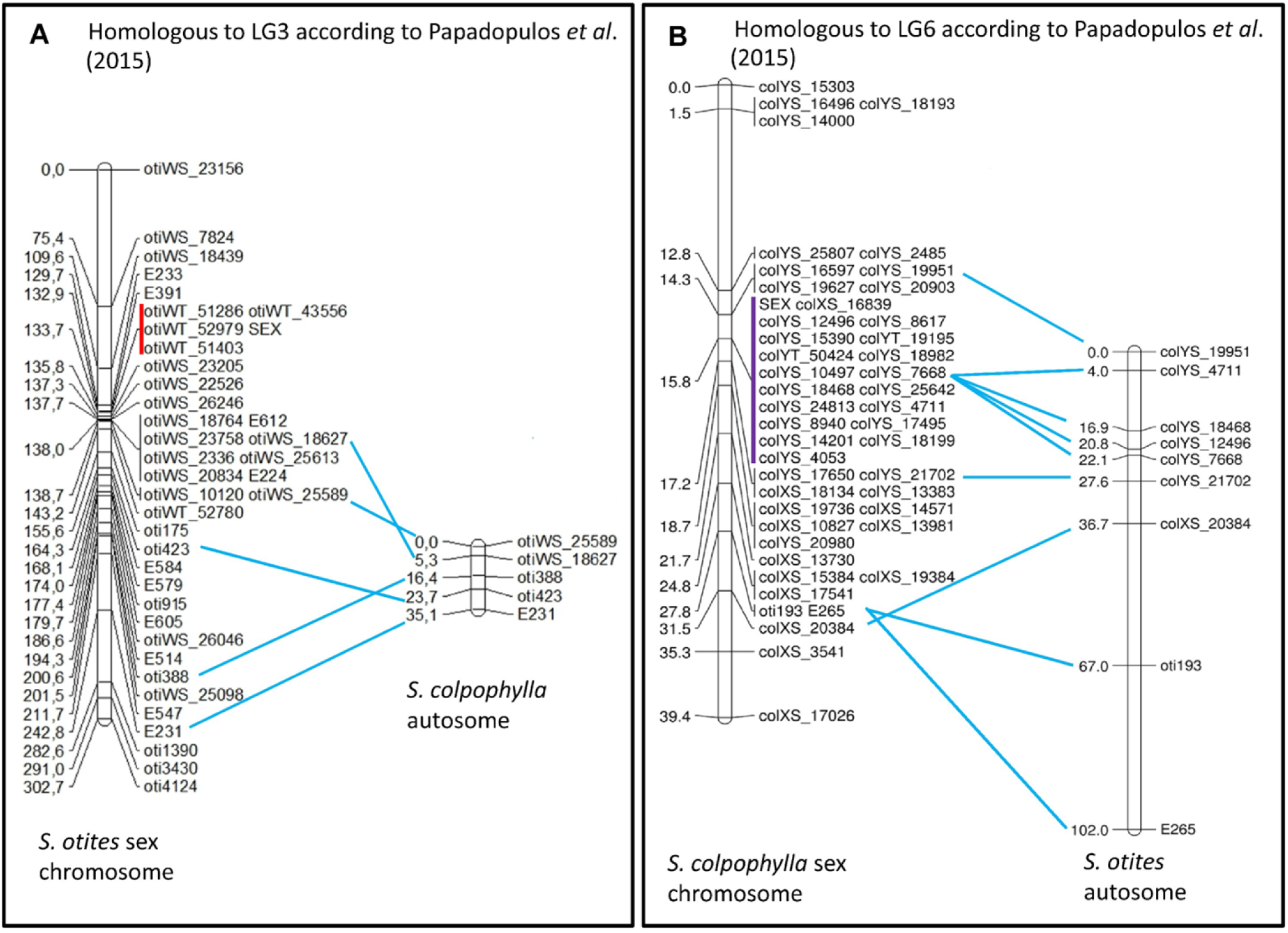
Comparison of the genetic maps of *S. otites* and *S. colpophylla*. The figure shows that the *S. colpophylla* homologs of the sequences that are sex-linked in *S. otites* are not sex-linked in *S. colpophylla* (a) and vice versa (b). The completely sex linked sequences (no recombinants found in the mapping population, and complete linkage disequilibrium in two natural populations) are marked by a red line (only in *S. otites*). The putatively completely sex linked sequences (no recombinant was found in mapping population but linkage disequilibrium in natural population was not tested) are marked by violet line (only in *S. colpophylla*).

#### S. otites

Candidate fully or partially sex-linked genes in *S. otites* were identified and genetically mapped (Fig. 2A) using several approaches (see Materials and Methods and Supporting information Table S1a). Initial 454 sequencing yielded 36 sequences suitable for genetic mapping in our *S. otites* family of 113 plants (see Material and Methods), which identified seven partially sex-linked sequences. These markers are denoted by names beginning with oti, followed by a number. Orthology between these pseudoautosomal genes identified the *S. otites* sex chromosome as the homolog of *S. latifolia* linkage group LG6, in the numbering of Bergero *et al.* ^28^, or LG3 in the numbering of Papadopulos *et al.*^27^. Ten further sequences from this *S. latifolia* linkage group were then identified and genetically mapped in our *S. otites* family; these markers’ names begin with E (according to the original *S. latifolia* sequences ^28^). As a second approach for discovering sex linked genes, Illumina RNAseq was performed for the parents of the same *S. otites* family and 6 progeny of each sex. This approach identified 318 sequences with potentially completely Z-linked and/or W-linked SNPs; these markers are denoted by names beginning with oti, followed by WS and a number (S denotes <u>S</u>NP and W means that the SNP was W-linked). 25 of the sequences identified as probably sex linked by analysis with the LINKYX^29^ pipeline (see Methods) were genetically mapped using SNP variants, of which 16 were identified as pseudoautosomal. In a third approach, we tested sequences that show female-specific transcription in leaves and flower buds (identified using LINKYX); this yielded a further set of 83 sequences detected only in females in both tissues (these markers are denoted by names beginning with otiWT, followed by a number, T means identified based on sex specific transcription). 29 sequences of this set of 83 sequences were further tested using PCR. This study identified one pseudoautosomal sequence (based on mapping an SNP in the mapping population) and four completely sex linked sequences (based on detecting a PCR presence/absence polymorphism in the mapping population). All four candidate fully sex-linked polymorphisms behaved as fully sex-linked based on complete agreement with the sex phenotype in two natural population samples; PCR amplifications of these sequences yielded products only from female genomic DNA samples, and not from males (Supporting information Table S2).

Overall, these three approaches yielded a map with a total of 4 fully sex linked and 34 pseudoautosomal genes in *S. otites*. Fig. 2A shows the genetic map based on all 38 markers (Supporting information Table S1A). The sex-determining locus maps at 138.0 cM, and only the four markers just mentioned showed complete sex linkage; almost the whole of the chromosome’s total map length of 302.7 cM is therefore pseudoautosomal, in two genetically large pseudo-autosomal regions.

To estimate the physical size of the fully sex-linked region, BACs carrying the four fully sex-linked sequences were isolated and sequenced. The W-linked BAC clones proved to have low gene density, consistent with this genome region being non-recombining, and their sequences revealed only one new sex linked gene (Supporting information Table S3). The sum of the lengths of the two smallest BACs sequences, plus the smallest distances of the mapped genes from the BAC ends in two another BACs, yields a size of at least 180 kb for the region.

#### S. colpophylla

In *S. colpophylla*, the LINKYX pipeline identified 88 sequences in our RNAseq data with SNPs suggesting either X- and/or Y-linkage. A further 1,671 sequences showed male-limited transcription in flower buds in the same sample of plants, but only 22 of these showed male-specific expression in leaves, and only two of them showed male-specific PCR amplification from genomic DNA (one is a sequence with no known homologues, and the other shows homology to F-box proteins). 19 sequences showed female-specific transcription in buds, but only one also did so in leaves, and no sequence showed female-specific PCR amplification from genomic DNA. SNPs in 48 of the 88 putatively sex-linked sequences were mapped in our mapping population of 69 plants. This confirmed 18 genes as fully, and 28 as partially, sex-linked (Supporting information Table S1B). In all 14 cases where homologs of sex-linked genes could be identified in either *S. otites* or *S. colpophylla*, sex-linked genes in one species are autosomal in the other (Fig. 2). This conclusion is further supported by screening homologs of sex-linked sequences from *S. otites* and *S. colpophylla* in recently mapped scaffolds from *S. latifolia*^27^. Using the numbering of *S. latifolia* linkage groups according to Papadopulos *et al.*^27^, 29 of the 38 *S. otites* sex-linked sequences map to the linkage group identified by our mapping, LG3 (see above), while nine map to five other LGs (Supporting information Fig. S1). 27 homologs of the 29 *S. colpophylla* sex-linked sequences map to LG6, and two to LG1 (using numbering by Papadopulos *et al.*^27^) (Supporting information Fig. S1).

Fig. 2B shows the genetic map of the *S. colpophylla* chromosome carrying the sex-determining locus. The genetic results are consistent with the expected male heterogamety, and our 45 markers place the estimated location of a male-determining locus at 15.8 cM, out of a total map length of 39.4 cM for this chromosome. Unlike *S. otites*, the fully sex linked region includes a high proportion of the sex linked genes identified. The 18 sex linked markers correspond to at a least 16 different genes (in two cases it is not possible to exclude the possibility that the mapped markers corresponds to distant parts of the same gene). These markers yielded no recombinants with the sex-determining locus in our family. However, as markers closely linked to the sex-determining region in *S. colpophylla* were preferentially selected for mapping, no reliable comparison can be made of the sizes of the non-recombining region with that in the other species studied here.

#### S. borysthenica

For *S. borysthenica*, no information on heterogamety was previously available. We tested sex linkage by LINKYX analysis of SNPs in Illumina RNAseq data from the parents and 12 progeny plants sampled from 32 progeny in our family from this species. This revealed 99 sequences showing probable Z- and/or W-linkage, 62 of the 318 sequences that showed Z- or W-linked SNP(s) in *S. otites* also appeared sex-linked in *S. borysthenica*, and 47 of the 99 sequences with evidence of sex-linkage in *S. borysthenica* were also sex-linked in *S. otites*. Therefore the same chromosome carries the sex-determining gene in both these species. 19 putatively sex-linked sequences from *S. borysthenica* showed homology with mapped *S. latifolia* scaffolds. Seven of these sequences show sex-linkage in both *S. otites* and *S. borysthenica*. Consistent with our identification of the chromosome carrying the *S. otites* sex-determining locus, 13 of them (7 newly identified in *S. borysthenica* and 6 which are sex linked both in *S. otites* and *S. borysthenica*) were in LG3 scaffolds, in the numbering of Papadopulos *et al.*^27^; with one exception, the scaffolds carrying homologs of the *S. otites* and *S. borysthenica* sex linked genes are in the upper arm of *S. latifolia* LG3 (Supporting information Fig. S1 and Fig. S2). Finally, we tested sex linkage of the *S. borysthenica* homolog of otiW43556, one of the four *S. otites* genes showing complete W linkage and female specific expression; for this gene, borW44973, no males showed W-linked alleles in our *S. borysthenica* mapping population of 32 plants.

### Phylogenetic analyses: Origin of female heterogamety within section *Otites*

Our Bayesian phylogenetic analysis yielded estimated phylogenetic relationships between members of the section *Otites* (see Fig. 1). *S. sibirica* appears to be a very close relative of section *Otites* (the estimated time of the most recent common ancestor is about 0.72 MYA). The dioecious section *Otites* must thus be very recent (estimated age is 0.55 MYA). *Silene sibirica* is gynodioecious, not dioecious, having female and hermaphrodite sex morphs. Gynomonoecious individuals with hermaphrodite and female flowers are also occasionally found (Supporting information Fig. S3).

The ancestor is inferred (see Methods) to have had female heterogamety in both the most recent common ancestor of the section *Otites*, and of the *Otites* group within this section, (Supporting information Table S4a,b). MultiState analysis in BayesTraits V3.0 evaluates this scenario as significantly more probable (5 times more, based on ratio of marginal likelihoods) than the next most probable scenario (positive evidence according to the scale proposed by Kass and Raftery^30^; positive evidence corresponds to values of 2×difference in ln(marginal likelihood) between 2 and 6). Given the tree, ancestral female heterogamety is consistent with the fact that both the extant species with female heterogamety (see Fig. 2) have sex-determining loci located on a chromosome homologous with the same *S. latifolia* LG (3 in the numbering of Papadopulos *et al.*^27^, 2015; see Supporting information Fig. S1).

Introgression is thought to occur from *S. colpophylla* into *S. pseudotites* (see the Discussion section), and therefore the male heterogamety shared by these two *Otites* group species could reflect gene flow from *S. colpophylla* into *S. pseudotites* (or its ancestor). We therefore also calculated the probabilities of all nine possible scenarios ((i) both section and group *Otites* ZW, (ii) section *Otites* ZW, group *Otites* XY, (iii) section *Otites* ZW, group *Otites* non-dioecious, (iv) section *Otites* XY, group *Otites* ZW, and (v) both section and group *Otites* XY), (vi) section *Otites* XY, group *Otites* non-dioecious), (vii) section *Otites* N, group *Otites* ZW), (viii) section *Otites* non-dioecious, group *Otites* XY) and (ix) both section and group *Otites* non-dioecious. We also allowed the origin of male heterogamety in *S. pseudotites* not to be an independent event (*S. pseudotites* could have either male or female heterogamety). The conclusion that the most recent common ancestor of both the section *Otites* and group *Otites* had female heterogamety is then 17 times more probable than the next alternative. As *S. colpophylla* males are heterogametic, this implies that there was a change in heterogamety, and also a change in the chromosome carrying the sex-linked genes, during the evolution of this species. The chronogram therefore suggests that male heterogamety evolved on the *S. colpophylla* branch within group *Otites* about 0.35 MYA (Fig. 1).

## Discussion

Our data confirm the male heterogamety in *S. colpophylla* and female heterogamety in *Silene otites* previously inferred using AFLP^18,31^ and markers from 454 sequencing^18^, and demonstrate the presence of a large region with highly suppressed XY recombination in *S. colpophylla* (corresponding to a large homologous region of at least 18 cM in *S. otites*), and our new data reveal that the chromosome carrying the sex-determining genes in *S. colpophylla* evolved from different a autosome pair from that carrying the *S. otites* sex-determining genes. The overall results (Fig. 1, Supporting information Table S4a,b) suggest that the ancestral state in both section *Otites* and group *Otites* was female heterogamety, and that the original sex determining region evolved on the homolog of autosomal linkage group 3 of *S. latifolia* (according to Papadopulos *et al.*^27^ numbering).

Male heterogamety in *S. colpophylla* and *S. pseudotites* could reflect evolution in a common ancestor of the group *Otites*. These two species probably share a common sex-determining locus as the sex chromosomes of *S. pseudotites* correspond to the LG6 in *S. latifolia* according to numbering by Papadopulos *et al.*^27^ (current results obtained by group of Prof. Pascal Touzet, University of Lille 1, Lille, France; Prof. Touzet personal communication). This fact suggests that males could have been the heterogametic sex in the common ancestor with *S. otites* (which must subsequently have changed to female heterogamety). This would have allowed time for the evolution of the large low recombination region in *S. colpophylla*. However, a large amount of time is not necessarily required, as a single large inversion could create such a region; to date, no evolutionary strata have been demonstrated in this species, so it is unclear whether multiple recombination suppression events have occurred (which would imply a large time). The possibility of a change from ancestral male heterogamety appears less probable than female heterogamety in the common ancestor of the group *Otites* (based on the analysis of the ancestral states, Supporting information Table S4).

A more plausible alternative that does not require such a change in *S. otites* is that the male heterogamety in *S. pseudotites* resulted from introgression of the XY sex determining system from *S. colpophylla*. The same localization and possible identity of the sex determining locus in *S. pseudotites* and *S. colpophylla* is in accordance with this view. Hybrids between *S. colpophylla* and *S. pseudotites*, and between *S. otites* and *S. pseudotites*, are thought to occur naturally in central eastern France, and hybrids between *S. colpophylla* and *S. otites* are phenotypically similar to *S. pseudotites*^32^. Our molecular data suggest that our *S. pseudotites* sample from Trieste could be a distinct species, sister to *S. otites* (we estimate the age of the most recent common ancestor to be > 0.2 MYA). Introgression from *S. colpophylla* cannot have occurred recently in Italian populations of *S. pseudotites* as there are no populations of *S. colpophylla* in Italy, but it could have occurred in the past. Introgression of a sex-determining region from *S. colpophylla* into *S. pseudotites* does, however, avoid the requirement that either ZW systems evolved twice independently from the same autosome (in both the *Otites* group species *S. otites* and the *Cyri* group species *S. borysthenica*), or that introgression of female heterogamety occurred from the *Cyri* group species (or their ancestor) into the *Otites* group species *S. otites*, which is improbable on geographic grounds. Species of the *Cyri* group have mostly eastern distributions (in Ukraine and Russia) or are endemic (*S. velebitica* in Croatia), while *S. otites* occupies western and central European localities^32^. Contact in the past cannot, however, be excluded.

In fish, many different autosomes have given rise to sex chromosomes in medaka and related fishes^33^, as we infer to have also occurred in *Silene*. An origin of *S. colpophylla’s* XY sex determining system from an ancestral ZW sex-determining system in section *Otites* is also consistent with the phenotypes of interspecific hybrids between *S. otites* and *S. pseudotites*^17,34^. Hybrids possessing both the W-chromosome and Y-chromosome have male phenotypes, i.e. the Y-linked sex determiner is dominant and epistatic to the previously existing factor. In the future, the study of synonymous site divergence between sequences of Y- versus X-linked genes, and W- versus Z-linked ones should enable us to further characterize the XY systems (in *S. colpophylla*, *S. pseudotites*) and the ZW systems (in *S. borysthenica*, *S. wolgensis* and *S. otites*).

Why might species in section *Otites* have changed heterogamety? The causes of turnovers in sex determination and switches in heterogamety have been classified into three main categories (reviewed by Beukeboom & Perrin^35^): (1) selectively neutral changes (2) changes driven by fitness differences between sex phenotypes and (3) sex ratio selection. Recently, modelling of finite populations has shown that genetic drift favors the spread and fixation of dominant masculinising or feminising alleles, which can lead to changed sex determination in a population in which a new genotype involved in sex determination (for example a dominant female-determining mutation on an X chromosome in an XY system) has no fitness advantage^36^; dominant alleles are therefore predicted to replace ones with lesser dominance. Changes in category 2 can be due to a new sex-determining mutation arising closely linked to a gene with a polymorphism for sexually antagonistic alleles (reviewed by van Doorn^37^, 2014), or may arise as a consequence of the degeneration of the original non-recombining chromosome (the mutational load hypothesis of Blaser *et al.*^38^). In this type of model, the combined effects of sexually antagonistic mutations and mutational load can potentially lead to repeated changes in sex determination among a limited set of chromosomes (the “hot-potato model“ of Blaser *et al.*^39^). In these models, the ancestral heterogamety is preserved after the changes, as is observed in anurans, though *Rana rugosa* is an exception^40,41^, and this is supported by the data from other vertebrates reviewed by Blaser *et al.*^39^. Finally, sex ratio selection can play a role if a population’s optimal sex ratio in the population is non-1:1, or if the sex ratio has been distorted by a genomic conflict (reviewed by Beukeboom & Perrin^35^).

All of these hypotheses require the new sex-determining gene to be epistatic to the original sex determining master gene (i.e. to determine sex even in the presence of the ancestral gene). Our results do not identify the mechanism responsible for the switch in heterogamety in section *Otites*. However, the male determiner of *S. pseudotites* is indeed epistatic to the female determiner of *S. otites*, as WY hybrid plants show the male phenotype^17^, as expected if female heterogamety was the primary sex determining system. As the age of the sex chromosomes in this section is very small, and our results suggest that the non-recombining region of the *S. otites* W-chromosome carries few genes, models involving genetic degeneration of this region of the W-chromosome appear improbable. The mutation load model is also unlikely, as changes in heterogamety have clearly occurred, in one direction or the other, and are not predicted by this type of model.

The extent of genetic degeneration is an important factor in determining whether a change in sex-determining locus also involves a change in the chromosome on which it is located, because such a change involves the pre-existing sex-determining chromosome becoming homozygous. If this chromosome carries a degenerated Y- or W-linked region, the change may be prevented due to low fitness or complete inviability of homozygotes of such chromosomes. Again, changed heterogamety in *Silene* section *Otites* is consistent with our observations suggesting that degeneration is likely to be minor in the ZW species. In such conditions, changed heterogamety is possible under the sexually antagonistic mutation model^42^. This model may explain the case of the sex determination switch in *Cichlidae*^43^, which also involved a change in the linkage group carrying the sex-determining region. We conclude that the change to a XY sex-determining system in section *Otites* could have involved either sexually antagonistic selection or neutral processes which allowed recruitment of a new epistatic sex-determining factor.

A difference in the ages, and the extent of genetic degeneration, can potentially explain the difference between the changes in section *Otites* species, versus maintenance of male heterogamety in species in section *Melandrium* (subgenus *Behenantha*), whose sex chromosomes have remained XY, and undergone no major changes apart from some translocations, including a reciprocal Y-autosome translocation in *S. diclinis* ^44^ and expansion of the pseudoautosomal region in *S. latifolia*^28^. The difference might reflect the very different extents of the sex-determining non-recombining regions. Species in section *Melandrium* have extensive non-recombining Y-linked regions, carrying large numbers of genes, including anther development genes essential for male fertility^45^–^48^ and genes essential for anther development^46^–^48^. Given its numerous genes, and the lethality of YY genotypes (reviewed by Westergaard^45^), the non-recombining regions of the *S. latifolia* Y-chromosome has probably undergone some degeneration, and there is direct evidence of this in *S. latifolia*^27,49^–^51^.

Interestingly, so-called “androhermaphrodites” (plants with male and hermaphrodite flowers) were found at an estimated frequency of 0.2% in two natural populations of *S. otites*^52^ (based on data of Correns). Their progeny included only males and androhermaphrodites, but no females^17,52^, suggesting, in combination with our current data, that their genotype was ZZ, not ZW. An alternative explanation could be that such the androhermaphrodites might possess W-chromosomes that are unable to suppress male function, and their female promoting function is compromised. In other words, it would mean that some W-chromosomes in some natural populations may be similar to Z-chromosomes, in terms of their sex-determining functions.

These findings suggest that the female-specific part of the *S. otites* W-chromosome is probably not essential for gynoecium formation, and that at least some Z-chromosomes lack no genetic information necessary for gynoecium formation. In speculating about the possible *de novo* evolution of ZW sex determination in section *Otites* species., the most parsimonious scenario is a two-locus gynodioecy model, as the very close relative (*S. sibirica*) is gynodioecious (a similar model may apply in the genus *Fragaria,* in which the same is true). If female heterogamety was the first sex determining system to evolve, male function must have been suppressed by a dominant male-suppressing mutation (rather than a recessive loss-of-function male-sterility mutation as proposed for the evolution of male heterogamety^1^. Such an effect could certainly arise by a dominant negative mutation directly creating a dominant female-determining chromosome (the mutation would define a proto-W-chromosome, see Fig. 3A). This mechanism is similar to the action of the W-linked homologue of the gene, which, in *Xenopus laevis*, counteracts DMRT1 expression by repressing the transcriptional targets of DMRT1^53^ (Yoshimoto *et al.*, 2008). Alternatively, it might involve a loss-of-function proto-W linked mutation of a haplo-insufficient gene important in anther development, such that proto-Z/proto-W plants, with only a single copy of the functional allele cannot develop normally functioning anthers (Fig. 3B). This mechanism resembles the presumed action of the DMRT1 gene in the gonadal sex determination in birds (reviewed by Kopp^54^).

**Figure 3.**
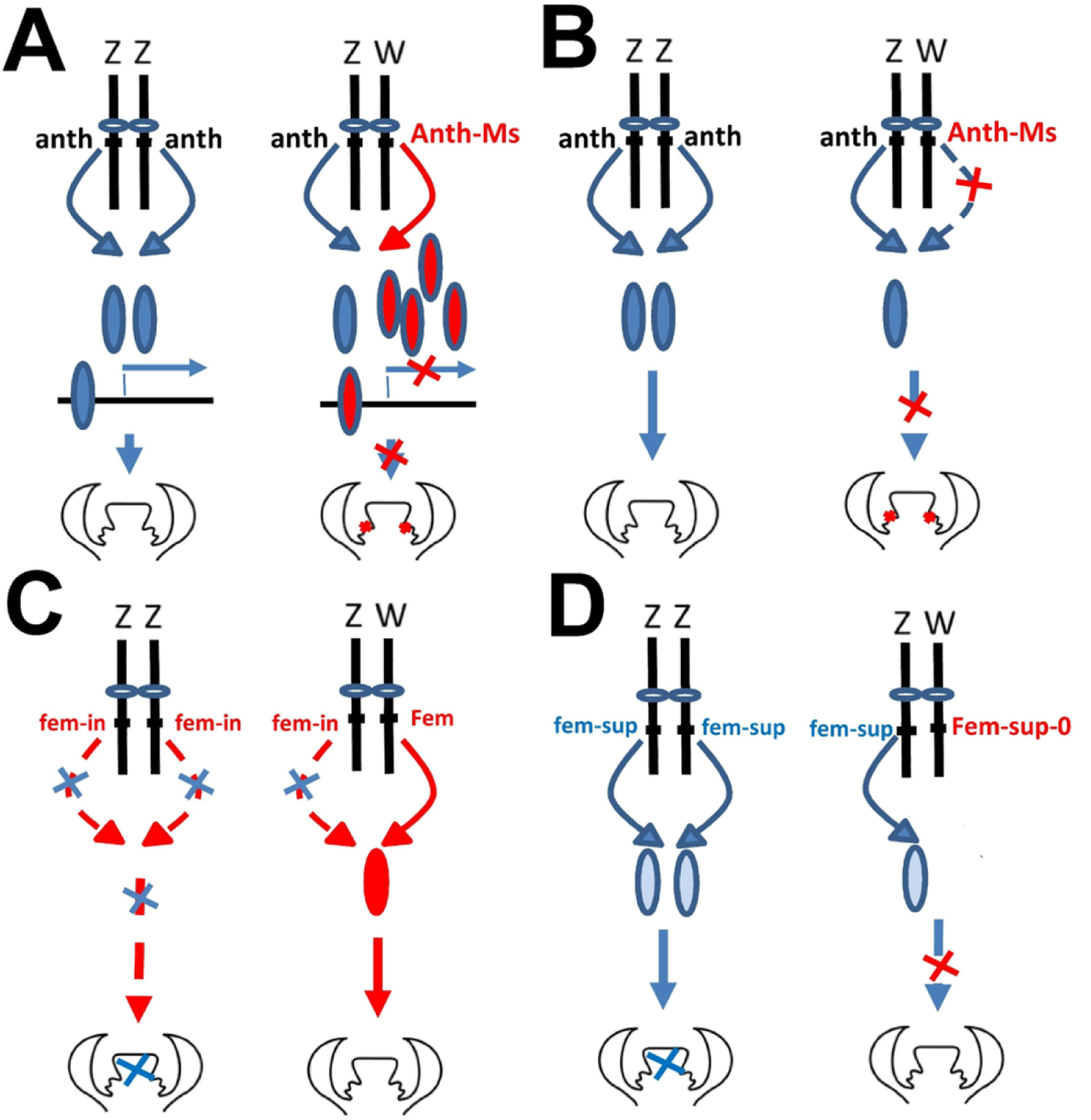
Possible routes to female heterogamety from gynodioecy in the section *Otites*. Possible types of anther suppression mutations on the proto-W-chromosomes (A, B) and possible ways of gynoecium suppression (C, D). **(A)** dominant negative effect based W-linked anther suppressor. Male sterility is caused by the repression of the transcriptional targets of the Z- linked transcription factor. W-linked inactive variant of a transcription factor competes the with the active variant in the linkage to its binding site in *S. otites.* (B) anther suppression caused by a haploinsufficient loss of function mutation on the proto-W-chromosome (C) A recessive loss-of-function mutation on the proto-Z-chromosome. The presence of androhermaphrodites in *S. otites* is hardly to explain in this case. (D) Recruitment of dosage dependent Z-linked allele involved in gynoecium suppression. No W-linked allele with such function is present. This hypothesis more parsimoniously explains the existence of ZZ androhermaphrodites. Note: Damage of anthers or gynoecium is indicated by red cross or blue cross symbols over their primordia.

To create males, a second mutation is required. A recessive loss-of-function female-sterility mutation (Fig. 3C) cannot readily account for the *S. otites* androhermaphrodites, which (as explained above) appear to be ZZ individuals. Therefore, it seems that one must invoke a dosage-dependent proto-Z linked allele causing gynoecium suppression when homozygous, but not when present in a single copy in ZW plants (Fig. 3D). As in the two-locus model of the origin of sex determination with male heterogamety (reviewed by Charlesworth^55^), this allele is more likely to invade the population if it occurs in a locus that is closely linked (in repulsion) to the male-suppressing locus. This dosage-based mechanism would again be similar to the proposed action of the DMRT1 gene in birds. The female-suppressing effect of DMRT1 in birds is, however, indirect through its role in the formation of male gonad from an originally bipotential tissue (reviewed by Kopp^54^) and so one master gene can perform both “male promoting” and “female suppressing” functions.

An interesting aspect of the scenarios outlined here, which assume dosage-dependence, is that the W-chromosome need not contain any master gene encoding a protein with a sex-determining function. This would make the ZW sex determining system prone to become epistatically recessive to the XY system. For example, a Y-chromosome might arise with a duplication of the Z-linked master gene. Under the model of Veller *et al.*^36^, no special driving force (such as a sexually antagonistic allele) is necessary, and the spread of the new dominant sex determining allele can occur by drift, potentially explaining the change(s) in section *Otites* within a short time period.

## Material and Methods

### Plant material and sex systems of the species studied

Information concerning all species included in the phylogenetic and/or genetic analyses is summarized in the Supporting information Table S5. Male heterogamety in *S. pseudotites* and female heterogamety in *S. wolgensis* were inferred from results of interspecific crosses^17,34^ and the current knowledge of the sex determining system in *S. otites* (Slancarova *et al.*^18^, and this work). *S. otites* was used as the second parent in the interspecific crosses by Newton^34^ and Sansome^17^. The male heterogamety has been recently confirmed in *S. pseudotites* by molecular data (current results obtained by group of Prof. Pascal Touzet, University of Lille 1, Lille, France; Prof. Touzet personal communication).

Candidate fully or partially sex-linked genes were identified using several different approaches (see Supporting information Table S1A,B). Sex linkage was tested in controlled crosses (F1 generation) in all three species studied; the family sizes were 113 (58 females and 55 males) in *S. otites*, 32 (15 females and 17 males) in *S. borysthenica*, and 69 (27 females and 42 males) in *S. colpophylla*. In *S. otites,* tests for sequences completely linked to the W-chromosome (showing presence/absence polymorphism in PCR reactions with primers targeting sequences for which preliminary data suggested sex linkage; presence of a product indicates linkage disequilibrium with a female-determining allele) were done by PCR amplifications in plants sampled from two natural populations from the Czech Republic, located in Rohatec (Hodonin district) (sampled 15 females and 17 males) and Brno-Kralovo Pole (Brno city) (sampled 12 females and 18 males).

### Illumina RNAseq sequencing, Roche 454 RNAseq sequencing and genetic analyses

Initial Roche 454 RNAseq of *S. otites*, using cDNA from the seed parent of the F1 population used by Slancarova *et al.*^18^ to identify the first sex-linked genes from this species, was performed by GATC Biotech (Constance, Germany). The assembly was performed using mira3 assembler^56^. Single copy sequences were chosen from genes in this dataset with homology to shared single copy nuclear genes in *Arabidopsis, Populus, Vitis* and *Oryza* (plant species of the dataset described by Duarte *et al.*^57^, to minimise amplification of paralogs, which would complicate tests for sex linkage. Tests for sex linkage, including partial sex linkage (of pseudoautosomal region, or PAR, genes) were done using the families described above, and identified several pseudoautosomal genes (see below). The linkage group carrying their *S. latifolia* homologs was determined using the *S. latifolia* mapping population of Bergero *et al.*^28^, and proved to be LG6 (in the numbering of Bergero et al.^28^). The *S. otites* homologs of other genes from this chromosome were then tested for sex linkage in *S. otites* and mapped using the same *S. otites* mapping population.

The main approach used in the manuscript both for the identification of sex linked sequences and for preparation of the sequence data for phylogenetic analysis was Illumina RNAseq. Illumina RNAseq (Illumina HiSeq 2000 PE, 2 × 50 bp and Illumina NextSeq 500 PE, 2 × 75 bp) sequencing experiments were performed at the EMBL Genomics Core Facility, Heidelberg (Germany). Sequencing libraries were prepared according to the Illumina protocol using oligo-dT attached magnetic beads. The sequences obtained in the present study are deposited in GenBank (www.ncbi.nlm.nih.gov/genbank/; see accession numbers in Supporting information Table S6. Adapters and low quality sequences were trimmed using Trimmomatic^58^, and errors corrected using Rcorrector^59^. An RNAseq dataset was assembled separately for each of the studied species using Trinity^60^. The longest ORFs were identified using Transdecoder and annotated using Trinotate^61^. 1082 orthogroups were used in phylogenetic analyses (see below), and were also annotated using the Blast2go pipeline^62^.

In *S. otites*, Illumina RNAseq sequences were obtained separately from both parent plants of the family used for genetic mapping (flower buds and leaves, separately), and from six progeny of each sex (flower buds only). For *S. colpophylla*, both flower buds and leaves were sampled from the seed parent of our family, but only leaves were available from the pollen donor. However, leaf and flower bud samples of a “full brother” of this plant were also sequenced, as well as from six progeny of each sex. In *S. borysthenica*, sequences were obtained from flower buds of both parents and a sample of their progeny (five males and seven females).

The Illumina RNAseq data were used for preliminary screening for candidate sex linked genes in *S. colpophylla*, *S. borysthenica* and *S. otites*. Sequences with sex linked inheritance patterns were first identified using the LINKYX pipeline^29^. This pipeline is based on preparation of the reference sequence using Trinity^60^ and the mapping of the reads to the reference using BWA^63^ (standard settings). Gene duplicates are removed using Picard’s MarkDuplicates function (http://broadinstitute.github.io/picard/). Subsequently, variants are called. SAMtools^64^ mpileup is set to -uf and bcftools view are used with following parameters: “-p 0.85 -cgv -”. For the identification of the sequences which are carrying sex-linked SNPs, the reference is assembled from reads of homogametic parent (XX mother or ZZ father) only to avoid possible problems caused by usage of sequence based on all individuals (for the details, see Michalovova et al.^29^). The rules for filtering based on SNPs are as described by Michalovova et al.^29^ (see also Supplementary Methods S1-S3). In contrast, the reference build from all individuals of the given sex is used for identication of the sequences that show sex specific expression. The rest of process is otherwise similar to the sex specific SNP detection. If the contigs are male-specifically expressed or are coming from the sequences that are present only in males the female reads are not mapping to them. The rules for filtering based on quantification are as described by Michalovova et al.^29^ (see also Supplementary Methods S4). To detect sequences with SNPs that have X-linked inheritance patterns in *S. colpophylla,* the pipeline was used without modifications. Modifications were made to reflect female heterogamety, in order to detect SNPs with Z- or W- linked inheritance patterns in *S. otites*, and Y-linked inheritance patterns in species with male heterogamety. In *S. borysthenica*, for which no previous genetic information was available, we tested for both male and female heterogamety. The LINKYX pipeline was also used to identify genes with sex-specific expression in *S. borysthenica*, *S. colpophylla* and *S. otites*. The latest version of the pipeline, including all the modifications used here, is available at GitHub repository: https://github.com/biomonika/linkyx. To validate the conclusions from LINKYX, reads from individual plants were mapped to the candidate sex-linked sequences using Bowtie 2^65^ and the output was visually checked in the IGV browser^66^.

The variants used for genetic mapping included DFLP (DNA fragment length polymorphism)^67^, intron size variants (ISVS, see Bergero *et al.*^25^), and single nucleotide polymorphisms (SNPs). SNPs in our mapping populations were genotyped as CAPS or dCAPS markers^68^, or by a high-resolution melting technique^69,70^. Genetic maps were constructed using JoinMap version 4.1^71^. We were using algorithm for outbreeder full-sib family (called CP as “cross pollinators”). The linkage groups were inferred based on independence LOD score higher than 4. We have used regression mapping and Haldane mapping function as usage of Kosambi mapping function is not supported for CP mapping populations in the JoinMap 4.1. Charts of genetic maps were produced using MapChart^72^. Homologies between the sex chromosomes of *S. otites*, *S. borysthenica* or *S. colpophylla* and *S. latifolia* chromosomes were further studied using blastn^73,74^ searches of *S. latifolia* scaffolds carrying homologs of genes showing sex linkage in section *Otites* species (minimal homology = 75%, minimal length of homologous region = 160 bp, and maximal E-value = 1 · e^-35^. For scaffolds that have been genetically mapped in *S. latifolia*^27^, we recorded the locations of those carrying the homologs of sex linked genes from section *Otites* species.

### Phylogenetic analyses – species tree estimation and analysis of ancestral states

Phylogenetic analyses were based on sequences of single-copy genes with orthologues that were found (based on Illumina RNAseqs and transcriptome assemblies) in 10 species of the subgenus *Silene* of the genus *Silene* (we refer to these as orthogroups). We identified 1,082 such orthogroups using Orthofinder^75^, which is based on an MCL algorithm^76^. For the phylogenetic analysis we mapped the reads from the particular plant samples to the reference sequence (each orthogroup was represented by the corresponding sequence from *S. otites)* using Bowtie 2^65^. We also applied a phasing procedure (to avoid biases in estimates of divergence times, as Lischer *et al*.^77^, 2014 have shown that unphased data can produce errors). We have performed read-based phasing for individuals without pedigree information and read-based pedigree phasing^78^ for samples with available pedigree information (*S. borysthenica*, *S. colpophylla* and *S. otites*), as implemented in the software WhatsHap^79^. Estimated haplotypes were extracted from VCF files using a custom python script and the BCFtools-call method of Samtools v1.3.1^80^. When several phasing alternatives existed, the most parsimonious (the one requiring fewest substitutions) were chosen. If more than one equally parsimonious phasing existed, the alignment was discarded from further analysis. Such parsimony phasing was applied to 40 alignments, and three were discarded due to phasing ambiguity. Low coverage sites with a read depth below 2 were masked using the BEDtools^81^ maskfasta method. Multiple sequence alignments were performed using the FFT-NS-i option^82^ of MAFFT v7.13^83^.

In order to facilitate a fully parameterized multiple coalescent analysis, only one species was used as the outgroup (*S. sibirica*, the closest non-dioecious species). To avoid possible paralogous sequences, all alignments with more than two alleles for any of the taxa were removed before further analyses. All remaining alignments were manually checked for phasing ambiguities, and branch lengths appearing excessive were excluded after visual inspection. Such grossly aberrant branch lengths were detected in one to a few alleles in fourteen alignments (2% of the dataset after removal of alignments with more than two alleles), which were discarded. In eight of these, these long branches appeared to be correlated with paralogy, i.e., the aberrant allele formed a separate branch from the other alleles, but the species topology was the same (termed ”mirror image topologies”). The final phased data set consisted of 662 alignments.

The data were analyzed using the Starbeast2 version 0.13 plugin^84^ and BEAST2^85^ version 2.4.7. We used a strict clock and a Jukes-Cantor substitution model for all alignments. A lognormal distribution with parameters (4.6, 2) was used for the speciation rate, and, for the population size, a lognormal distribution (-7, 2) was used as prior. The Markov Chain Monte Carlo was run for 1,000,000,000 iterations. For the time estimates, we used calibration based on estimated divergence per synonymous site *dS* (substitutions per synonymous site) and the divergence time estimates for several angiosperm lineages (*Brassicaceae*, *Malvaceae*, *Euphorbiaceae*, *Fabaceae*, *Cucurbitaceae*, *Rosaceae*, *Solanaceae*, and *Poaceae*), so as not to depend on a single fossil record or phylogenetic tree position, which yielded a mean substitution rate of 5.35 × 10^−9^ synonymous substitutions/site/year^86^. Mean *dS* values between the *S. sibirica* outgroup species and each section *Otites* species were estimated for each gene using the HyPhy package^87^. The mean *dS* is 0.0077; using the above angiosperm substitution rate per year, and assuming one generation per year for the Silene species, the estimated age of the most recent common ancestor of the *S. sibirica* and the section *Otites* species is about 720 thousand years (SE = 13 thousand years).

Ancestral state analysis was performed in BayesTraits V3.0 using the MultiState method^88^. The analysis took into account all currently available information about male and female heterogamety in the section *Otites*, including previously published phenotypic data^17,34^. All nine possible combinations of ancestral states of the most recent common ancestor of the whole section *Otites* and of the group Otites (see Supporting information Table S4a,b) were tested ((i) section *Otites* ZW, group *Otites* ZW, (ii) section *Otites* ZW, group *Otites* XY, (iii) section *Otites* ZW, group *Otites* N, (iv) section *Otites* XY, group *Otites* ZW, and (v) section *Otites* XY, group *Otites* XY), (vi) section *Otites* XY, group *Otites* N), (vii) section *Otites* N, group *Otites* ZW), (viii) section *Otites* N, group *Otites* XY) and (ix) section *Otites* N, group *Otites* N; female heterogamety-ZW, male heterogamety –XY, non-dioecious-N). As the input we used the set of the species trees obtained using the Starbeast2 plugin^84^ and BEAST2^85^. The chains were run for 10^9^ states, using a gamma hyper-prior (mean from uniform: 0-10, variance from uniform: 0-10). Again, the process was checked using Tracer^89^. The marginal likelihoods for the Bayes factors were obtained using the stepping-stones method following the BayesTraits V3.0 tutorial. The significance of differences was evaluated according to Kass and Raftery^30^.

### BAC library construction, screening and the subsequent analyses

A partial genomic BAC library was constructed for *Silene otites* (sample NN1 from the Nantes (France) population) using high molecular weight DNA prepared from flow sorted nuclei. The protocol was modified from Cegan *et al.*^90^, for the details see Supporting information Methods S5. Library screening was performed by radioactive hybridization with ^α^32P using the Prime-It II Random Primer Labelling Kit (Stratagene) following Cegan *et al.*^91^. Probes were prepared by PCR amplification of selected markers (see below). Positively hybridizing BAC clones were selected, and the presence of a gene of interest was confirmed by PCR and sequencing. BAC DNA was isolated from the positive BACs and sequenced using an Illumina MiSeq (2×300PE) machine at the Centre of Plant Structural and Functional Genomics (Olomouc, Czech Republic). The reads were assembled *de novo* using the Edena assembler^92^.

## Acknowledgements

This project was supported by a grant from the Czech Science Foundation (GACR) 13-06264S to BJ. Computational resources were provided by the CESNET LM2015042 and the CERIT Scientific Cloud LM2015085, provided under the programme “Projects of Large Research, Development, and Innovations Infrastructures”. We thank Dr. Jiri Danihelka (Masaryk University, Brno) and the Botanical Garden of Trieste for the seed material. We also thank to Monika Cechova (Pennsylvania State University), Pascal Touzet (University of Lille1, Lille) and Gabriel Marais (University Lyon 1, Lyon) for discussions.

## Author Contributions

BJ, VB, RG, JS, RH and RB conceived and designed the experiments. VB, RG, JS, BJ, and RB performed the experiments. BJ, RC, PC, BO, JZ, VB, RG, and VK analysed the data. BJ contributed reagents/materials/analysis tools. BV, VB, RG and RH discussed the paper, and BJ, dJZ, DC and BO wrote the paper.

## Additional Information

Supplementary information accompanies this paper at https://will be completed in the final version

## Data availability

The Illumina RNAseq raw data and Roche 454 RNAseq raw data were deposited to GenBank (www.ncbi.nlm.nih.gov/genbank/; see accession numbers in Supporting information Table S6).

